# Larval surveys reveal breeding site preferences of malaria vector *Anopheles* spp. in Zanzibar City

**DOI:** 10.1101/2024.10.23.619825

**Authors:** Kaeden K. Hill, Dickson Kobe, Katharina Kreppel

## Abstract

Malaria is a serious illness that causes over 500,000 deaths annually worldwide, mainly affecting children under five years of age. Tanzania, in Africa, contributes to approximately 4% of those deaths. Malaria is caused by *Plasmodium* spp. parasites, and it is vectored by mosquitoes in the *Anopheles* genus. In Zanzibar City, the capital of the Zanzibar archipelago in Tanzania, the incidence of malaria has decreased over the past few decades due to standardized treatment protocols in hospitals and dispensaries and public health interventions targeting adult mosquitoes. However, the incidence remains between 1-2%, with an increasing trend observed over the past few years likely stemming from continued exchange of *Plasmodium* spp. from other malaria-endemic areas (including mainland Tanzania). While larvicides are a powerful tool to reduce vector populations, this control strategy relies on information on breeding sites and their productivity. In Zanzibar City, no larval surveys have been done in the last few years. Our aim was to characterize *Anopheles* spp. breeding sites in Zanzibar City during the rainy season. We first conducted systematic larval surveys across 16 semipermanent/permanent water bodies and 30 temporary water bodies. Then, we used Principal Component Analysis and logistic regression to model the effect of physical and chemical parameters, and rainfall on *Anopheles* presence. We found that *Anopheles* spp. utilize mostly concrete, semipermanent breeding sites with high levels of dissolved oxygen concentrations but can utilize natural sites after heavy rains. The logistic regression model incorporating rainfall was able to predict the presence of Anopheles larvae with a positive predictive power of 65.7% and a negative predictive power of 88.8%. The data from our study suggest that *Anopheles* spp. have not yet expanded to using more polluted breeding sites in Zanzibar City (as they have in some mainland locations). These results can inform targeted larvicidal strategies in Zanzibar City.

## INTRODUCTION

Malaria is a vector-borne disease that causes over 500,000 deaths annually [1]. Malaria is caused by the *Plasmodium* parasite with a life cycle involving stages in the human host and in the female *Anopheles* spp. mosquito vector. Throughout Tanzania (including the Zanzibar archipelago), malaria is one of the most widely-spread mosquito-borne diseases, with at least 2.5 million cases reported in 2022 [1–3]. Tanzania has contributed 4% of the total malaria deaths worldwide [1].

Understanding the local ecology of the *Anopheles* mosquitoes is crucial to inform control strategies. Amid changing climate conditions and an increase in tourist traffic, Zanzibar City - the largest city of the Zanzibar archipelago that includes Stone Town (a UNESCO World Heritage Site [4]) - has experienced a recent increase in malaria cases. In 2009, the Zanzibar Ministry of Health initiated a set of policies to eliminate malaria on the archipelago, including increased indoor spraying, a systematic distribution of insecticidal nets, and a surveillance and case-detection system [5,6]. Zanzibar has therefore been able to reduce malaria transmission between 2005 and 2015 [7].

However, malaria incidence rate of 2.7 cases per 1,000 people in 2017 has increased to 3.6 in 2021 [6,8]. The increase in cases is thought to be due to an expansion of tourism and a greater influx of people from mainland Tanzania potentially bringing the infection with them [9,10]. This situation suggests that entomological interventions may be most effective in eliminating malaria, as it is the resident population of the vector mosquitoes that allow for the imported *Plasmodium* parasites to be transmitted. Due to their low mobility and fully-aquatic nature, mosquito larvae that develop in freshwater bodies are effective targets for reducing the abundance of adult vector mosquitoes and overall malaria transmission [11]. Therefore, the aim of this study was to determine which type of breeding sites are used by *Anopheles* mosquitoes during the rainy season in Zanzibar City and what characterizes them by identifying the physicochemical parameters.

While urbanization was originally thought to decrease malaria transmission, instances of unplanned, rapid urbanization in sub-Saharan Africa put the population at risk of malaria [12–15]. Zanzibar City has expanded 3.8% per year since 2004, leading to unplanned urbanization with poor integration of proper water drainage into the city infrastructure potentially increasing mosquito breeding sites [16]. Therefore, urbanization has likely contributed to enhanced malaria transmission in recent years [17]. With a population of 800,010 people in 2023, and a population growth rate of 4.48%, this trend is likely to continue [18]. The annual rainy season between April and May leads to pooling of stagnant freshwater in the urban environment and thus contributes to greater numbers of mosquitoes and potential disease vectors. Despite existing drainage of the streets of Zanzibar City, excess water runoff can lead to the expansion or flooding of natural water pools. Zanzibar City, as a tourist destination, features aesthetic ponds and nonfunctional fountains that often contain stagnant water, creating breeding sites for mosquitoes (particularly within historic Stone Town).

Additionally, climate change has resulted in shorter but heavier rains during the March-May rainy season, likely increasing the number of potential urban mosquito breeding sites in Zanzibar City [19,20].

Currently, larvicidal strategies are rarely used in Africa [21,22]. However, because *Anopheles* mosquitoes on Unguja (the largest island of the Zanzibar archipelago where Zanzibar City is located) commonly bite humans outdoors, show both zoophilic and anthropophilic behaviors, and are becoming increasingly resistant to pyrethroid insecticides [23], larvicidal strategies may be an effective complement to existing insecticidal measures.

For larvicidal strategies to be effective against *Anopheles* mosquitoes, the defining characteristics of *Anopheles* breeding sites must be known. However, *Anopheles* breeding habits vary by geographical location and season, making it difficult to use studies conducted in other areas or at other times [24–26]. To our knowledge, no preexisting data regarding *Anopheles* breeding sites in Zanzibar City is publicly available. We set out to address the lack of data by conducting larval surveys and breeding site characterizations for *Anopheles* larvae in Zanzibar City during the rainy season. We found consistently high *Anopheles* larvae abundance in concrete, semipermanent structures with high dissolved oxygen levels, but also high *Anopheles* larvae abundance in natural semipermanent areas with high dissolved oxygen levels after heavier rains. Together, our data suggest that *Anopheles* mosquitoes use predictable breeding sites in Zanzibar City, supporting the current targetability of *Anopheles* mosquito larvae as a strategy to prevent malaria transmission.

## METHODS

### Study Site

Zanzibar City is a popular tourist destination located on the Western coast of Unguja, the largest island of the Zanzibar archipelago (1,666 km^2^) off the coast of East Africa. Zanzibar city is located at approximately 6°10’ S 39°12’ E. The climate of Zanzibar City is tropical, with an average annual rainfall of approximately 1,521mm [27]. Zanzibar City (and the entire Zanzibar archipelago) experiences two rainy seasons: short rains in October-December and long rains from March-May. The elevation of Zanzibar City ranges from 2m to approximately 45m above sea-level. As of 2023, Zanzibar City has a population of 800,010 people, which represents approximately 44% of the total population of the Zanzibar archipelago (1.8076 million people) [28].

### Sampling Locations

Between April 11, 2024, and May 1, 2024, 16 permanent or semipermanent sites and 30 temporary sites were sampled for *Anopheles* mosquito larvae (**Fig. 1**). Sites were initially found through reconnaissance or asking community members during the first week of the study. Because only publicly available water bodies were surveyed (and few such water bodies were found in the city’s Eastern sprawl), most study sites were near the more developed Stone Town and the densely populated neighborhoods nearby. Semipermanent sites were defined as permanent structures that are occasionally drained (but rarely during the rainy season), while permanent sites were defined as permanent structures that contain water continuously throughout the year.

**Figure 1:**
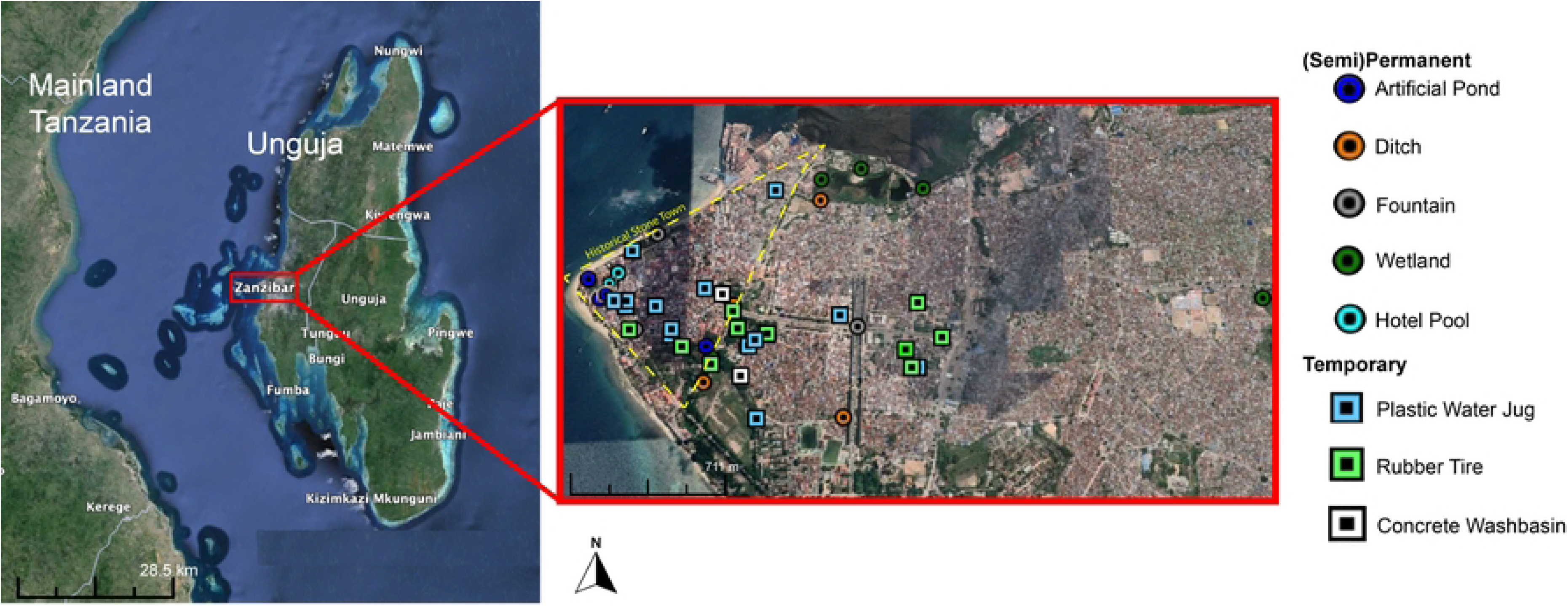
Map of Zanzibar City, Tanzania, showing sites sampled for mosquito larvae.

Semipermanent and permanent sites sampled included unfinished or nonfunctional fountains (n=3), artificial ponds (n=4), roadside ditches (n=3), and wetlands (n=5) (**Supplementary** Figure 1). Temporary sites included tires (n=10), plastic water containers (n=18), and small concrete wash basins (n=2). Each permanent or semipermanent site larger than 4m in perimeter was divided into semi-randomized 0.5m x 0.5m subsites. a random number was generated that indicates the 0.5m x 0.5m site to sample. Before sampling, however, the 0.5m x 0.5m subsite was inspected to ensure that it could physically house mosquito larvae (e.g. adequate, stagnant water present), and that it was accessible with a larval dipper. If a randomly generated subsite could not be sampled, the nearest possible subsite was sampled instead. Once the subsite was identified, a 0.5m x 0.5m quadrat was placed into the water. At each site, data was collected at five subsites unless the site was smaller than five 0.5m x 0.5m quadrats (referred to herein as “subsite”). Because mosquito larvae distribution is not evenly distributed across larger habitats [29], each subsite was treated independently during analysis. Notably, all *Anopheles* species on Unguja are capable of transmitting *P. falciparum* [10,30], so mosquito larvae were identified to genus level. Additional mosquito genera on Unguja include *Culex* and *Aedes* [31].

### Collection and Identification of Mosquito Larvae

Mosquito larvae were sampled using a 500mL or 80mL dipper (depending on the size of the water body) and a white inspection tray. Five dips were conducted per 0.5m x 0.5m subsite, and the percentage of dips containing larvae of each mosquito genus was calculated. Dipping was performed by the same individual at all sites and visits, except one visit to two wetlands and one drain (no *Anopheles* larvae were found at those sites during either visit with either field technician). Other data collected for each dip included the number of mosquito larvae of each genus found per dip, and the number and name of other macroinvertebrates. Predators of mosquito larvae at the dipping site were also recorded.

Mosquito larvae were identified to the genus-level using a hand lens and identification keys [32,33]. Pupae were not recorded due to difficult identification. If a particular larva could not be identified using a hand lens, the specimen was transferred to the laboratory for examination using an XSZ-107BN Series light microscope.

### Measurements of Physical Parameters

All physical parameters were measured within each 0.5m x 0.5m subsite before larvae were sampled. Coordinates were taken using the iPhone 11 compass. Outdoor and water temperature were both measured using a standard glass lab thermometer. Percentage vegetation cover was estimated by eye and included all living plants emerging at the surface of the subsite (e.g. patches of grass emerging from the water in a shallower subsite). Depth was measured within the subsite using either a metal measuring tape or a measuring tape with a weighted end. Other binary (yes or no) physical parameters recorded were: trash within 3m of the subsite, large dump (> 10 pieces of trash) within 3m of the subsite, semipermanence of dipping site, and whether the dipping site was constructed out of concrete, plastic/rubber, or natural. Additionally, the perimeter was estimated and placed into one of five categories (1= <1m, 2=1-5m, 3=5-10m, 4=10-20m, 5= >20m). All daily precipitation data was obtained from the Visual Crossing website (accessed April 27, 2024), which compiles data collected at the HTZA weather station (-6.22, 39.22) and the Abeid Amani Karume International Airport weather station (-6.22, 39.23) approximately 5km from Zanzibar City [34].

### Measurements of Chemical Parameters

Dissolved oxygen concentration, dissolved oxygen saturation, pH, relative turbidity, and relative chlorophyll content (estimated by relative intensity of green pigment in the water) were measured in each subsite on at least one of the visits for each site. Dissolved oxygen was measured on-site using a Ecosense DO200A within each 0.5m x 0.5m subsite before larvae were sampled. The instrument was calibrated before the first use using the manufacturer’s protocol. Both dissolved oxygen in parts per million (ppm) and percentage saturation were recorded. For the remaining chemical parameters, a water sample was collected in a cup (after first rinsing the cup in water from that subsite) and transported to the laboratory for immediate analysis. pH was measured using a Hanna HI991001 pH probe. In between measurements, the probe was briefly rinsed in deionized (DI) water and wiped off. The pH meter was calibrated before the first use using the manufacturer’s protocol. Salinity was measured using a MarineDepot refractometer that was calibrated daily using DI water. Values for relative turbidity and chlorophyll content were obtained by taking photos of the water inside a clear cup using an iPhone 11 camera (with flash) and analyzing the color distribution using the color histogram function on Fiji ImageJ2 [35] (2.14.0). The type of cup and the white background were kept constant across all photos. The mean color intensity for the red, green, and blue color channels in a selected area inside the cup were recorded and normalized to the mean values for a sample of DI water. The relative turbidity was therefore calculated by adding the normalized mean intensities for the red, green, and blue channels together, and subtracting that value from three (since a total normalized color value of three indicates that the red, green, and blue channel intensities were identical to those of DI water and is therefore the maximum whiteness/clarity for water). Relative chlorophyll content was estimated using the ratio of normalized mean green intensity to normalized mean blue intensity, as this value captures both brighter greens (DI-like normalized green intensity, lower normalized blue intensity), and darker greens (lower normalized blue and green intensity).

### Data Analysis and Statistics

All statistical analysis was done using GraphPad Prism 10 (10.1.1). The Anderson-Darling test was used to test for normality, and either Mann-Whitney tests or Kruskal-Wallis tests with uncorrected Dunn stepdown tests were used to compare groups of unpaired non-parametric data. As each pairwise comparison stood alone, no corrections were used. All non-parametric, paired data was analyzed using a paired Wilcoxon test. All parametric, unpaired data was analyzed using an unpaired Student’s T-test with Welch’s correction. All categorical data was analyzed using either a Fisher’s Exact test (for contingency tables with any frequencies below 10) or a Chi-Square test for independence. Groups of normally distributed data were analyzed using unpaired Student’s T-tests. Principal component analysis was conducted to determine which parameters to further investigate using data standardized to a mean of zero and standard deviation of one. Principal components were selected using parallel analysis.

While principal component regression (PCAR) was run on all physical predictors included in the study, a multiple logistic regression model was built using forward selection of parameters that were significant in the PCAR and chemical/interaction parameters for which there was evidence of a relationship. The interaction terms included in the multiple logistic regression model test whether concentration dissolved oxygen affects the presence of *Anopheles* differently if the subsite was semipermanent, made from concrete, or if rainfall levels were higher, and whether rain within 48hrs affects the presence of *Anopheles* differently in concrete or in natural subsites. A significance level of 0.05 was used for all statistical tests. All maps were generated using Google Earth Pro (7.3.6.9796).

### Statement of Ethics

Ethical approval for this study was granted by the World Learning Inc. Institutional Review Board, and a student research permit was granted by the Revolutionary Government of Zanzibar. Although most land accessed was publicly owned, permission to sample was obtained from the relevant owners whenever a water body crossed into private property. No protected species were sampled.

## RESULTS

### Characterization of physical properties of *Anopheles* breeding sites

To test whether *Anopheles* prefer permanent/semipermanent sites or temporary sites in Zanzibar City during the rainy season, we compared the number of sites with *Anopheles* larvae between permanent/semipermanent and temporary sites. We found that 56.25% of the permanent or semipermanent sites contained *Anopheles* larvae during at least one of the times the site was sampled, while only 10% of the temporary sites contained *Anopheles* larvae (Fisher’s Exact Test, *P*=0.0013, OR = 11.57, 95% CI = 2.359-44.85) (**Supplementary Table 1**). The three temporary sites with *Anopheles* larvae included two plastic water containers and one rubber. The permanent or semipermanent *Anopheles* breeding sites were in both Stone Town and the greater Zanzibar City area, while the temporary *Anopheles* breeding sites were only found in the Zanzibar City area outside of Stone Town (**Fig. 2A**). Although not included in the formal analysis because pools are subject to more potentially confounding anthropogenic variables, three pool drains across two hotel pools in Stone Town were sampled for mosquito larvae, and no larvae were found. Additionally, six large rain tanks for drinking water were visually inspected for larvae (but not sampled due to ethical issues surrounding conducting larval dips in drinking water). Among the three tanks inspected that were left open, two had *Culex* larvae present. None of the three tanks inspected that were closed had any larvae.

**Figure 2:**
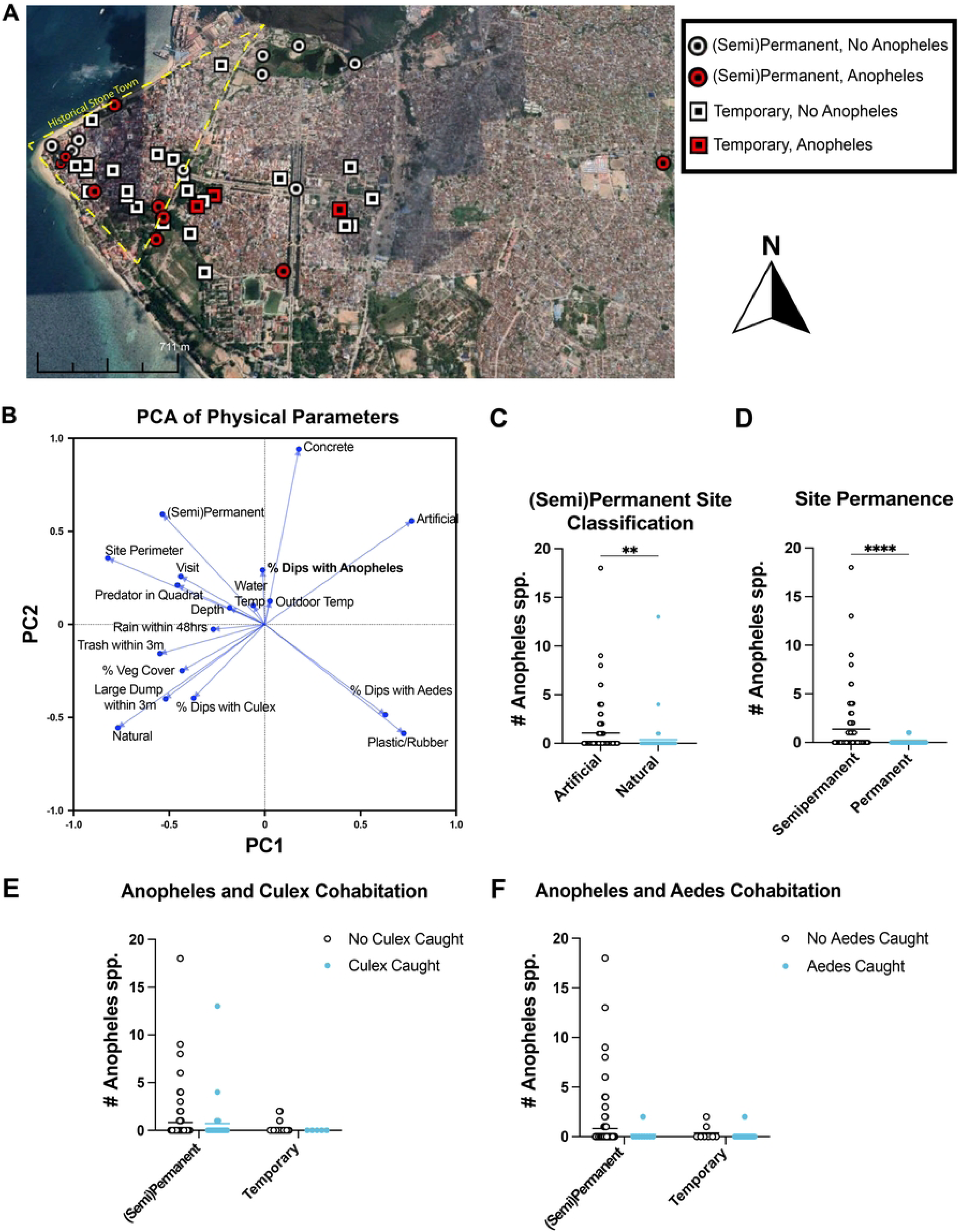
**Characterization of physical qualities of *Anopheles* breeding sites in Zanzibar City, Tanzania**. **A**) Map showing site permanence and whether they contained *Anopheles* larvae in either visit. **B**) Principal component analysis graph showing loadings of physical parameters. **C**) Distribution of *Anopheles* larvae abundance in artificial and natural subsites. **D**) Distribution of *Anopheles* larvae abundance in semipermanent and permanent subsites. **E**) Distribution of *Anopheles* larvae abundance in subsites of different site types in the presence or absence of *Culex* larvae. **F**) Distribution of *Anopheles* larvae abundance in subsites of different site types in the presence or absence of *Aedes* larvae. Statistical significance determined using a Mann-Whitney Test. For all pairwise comparisons: * *P*<0.05, \*\**P*<0.01, \*\*\**P*<0.001, \*\*\*\**P*<0.0001.

The percentage of dips at each subset with *Anopheles* was associated with concrete, artificial, and permanent/semipermenent subsites, in addition to water temperature and outdoor temperature (**Fig. 2B**). The percentage of dips with *Culex* was associated with natural subsites and the presence of visible trash, while the percentage of dips with *Aedes* was strongly associated with subsites made of plastic or rubber.

There was also statistical evidence for the permanent/semipermanent artificial subsites harboring more *Anopheles* larvae than permanent/semipermanent natural subsites (Two-tailed Mann-Whitney U test, *P*=0.0022, U_(93, 54)_ = 1953) (**Fig. 2C**). Significantly more *Anopheles* larvae were caught at semipermanent subsites (Two-tailed Mann-Whitney U test, *P*<0.0001, U_(84, 63)_ = 1701) (**Fig. 2D**). Although cohabitation with other mosquito larvae genera was uncommon, we found that there was no significant difference in abundance of *Anopheles* larvae in subsites with either *Culex* or *Aedes* larvae present versus subsites without *Culex* or *Aedes* larvae present (**Fig. 2E-F**). Last, we used principal component analysis regression (PCAR) to estimate the predictive capability of the parameters included in the PCA analysis (**Supplementary Table 2**).

Parameters with significant positive predictive capabilities included permanence/semipermanence, artificial, and concrete. Parameters with a significant negative predictive capability included percentage of dips per subsite with *Culex*, percentage of dips per subsite with *Aedes*, vegetation cover, the presence of a large dump within 3m, plastic/rubber, and natural. While the intercept was nonsignificant, the overall PCA regression was significant by analysis of variance (F_(3, 173)_ = 3.288, *P*= 0.0221).

### Characterization of chemical parameters of *Anopheles* breeding sites

We next analyzed which chemical parameters were associated with the presence of mosquito larvae of each genus. The percentage of dips per subsite with *Anopheles* was associated with dissolved oxygen concentration, dissolved oxygen saturation, and pH. The percentage of dips per subsite with *Culex* was strongly associated with salinity, and the percentage of dips per subsite with *Aedes* was associated with relative turbidity and estimated chlorophyll content (**Fig. 3A**). Although the distribution of the number of *Anopheles*, *Culex*, or *Aedes* larvae caught plotted against dissolved oxygen levels was not normally distributed (Anderson-Darling Test, *P*<0.0001), Gaussian curves were used to visualize which dissolved oxygen levels were associated with a higher larval abundance for each genus in permanent/semipermanent subsites. A least squares regression comparison of the Gaussian fits of larvae abundance plotted against dissolved oxygen saturation/concentration suggested that distributions differ between genera (Extra sum of squares F test, F_(6, 293)_ = 6.897, *P*<0.0001). The distribution of *Anopheles* larvae abundance was shifted towards higher dissolved oxygen levels (both concentration and saturation) than the distribution of *Culex* larvae abundance, while *Aedes* larvae were found in low abundance across a wide range of dissolved oxygen levels (**Fig. 3B-C**). While a Gaussian curve could not be fit for larvae abundance plotted against pH, the distribution of *Anopheles* larvae abundance was shifted towards higher pH levels than *Culex* (**Fig. 3C**). *Aedes* larvae were found in low abundance across a wide range of pH levels. Turbidity appeared to have little effect on the distribution of larvae abundance from different genera (**Fig. 3E**).

**Figure 3:**
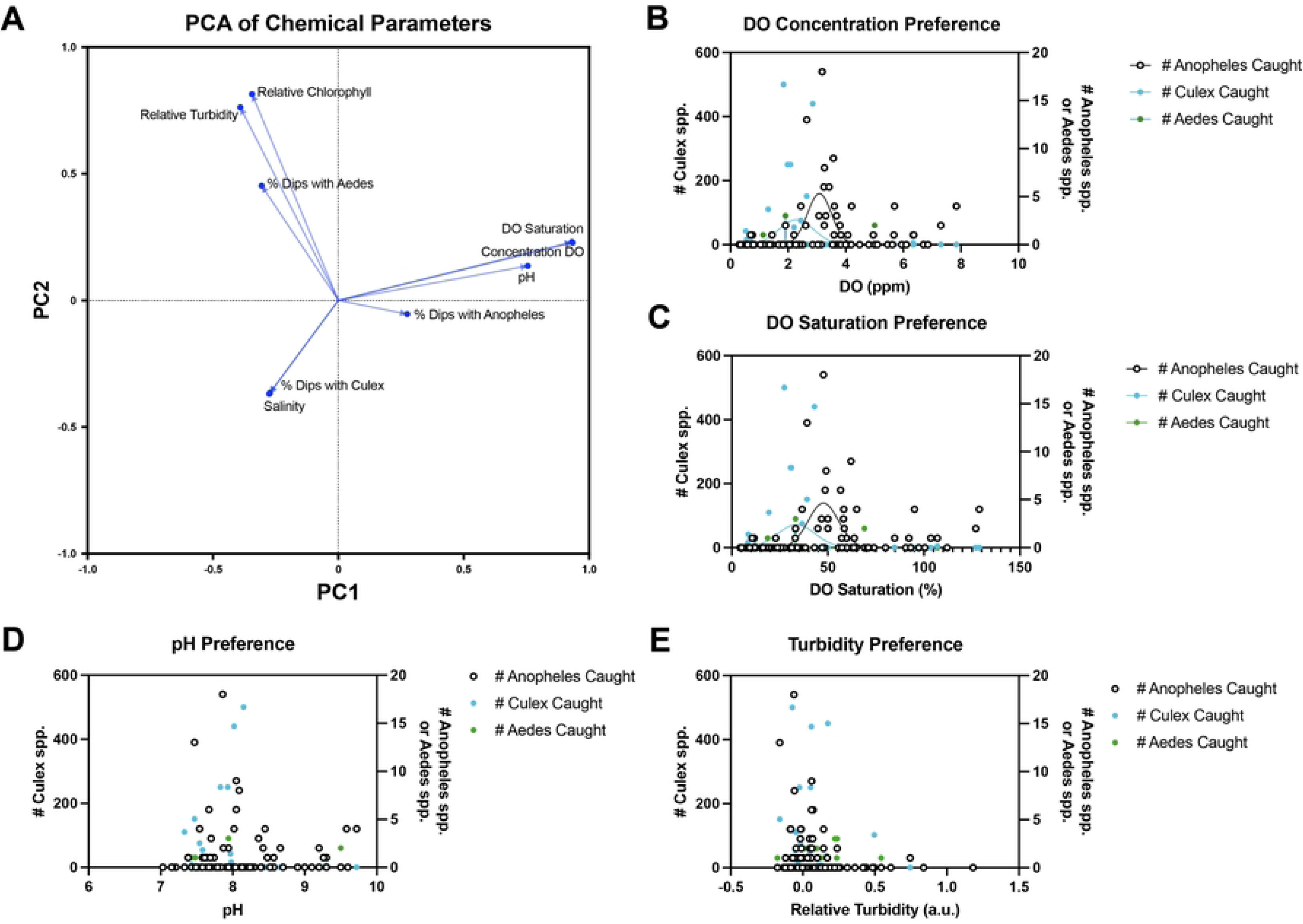
Characterization of chemical parameters of *Anopheles* breeding sites in Zanzibar City, Tanzania. A) Principal component analysis showing loadings of chemical parameters. **B**) Distribution of *Culex* abundance (left axis) and *Anopheles*/*Aedes* abundance (right axis) at each subsite plotted against dissolved oxygen concentrations. **C**) Distribution of *Culex* abundance (left axis) and *Anopheles*/*Aedes* abundance (right axis) at each subsite plotted against percentages of dissolved oxygen saturation. **D**) Distribution of *Culex* abundance (left axis) and *Anopheles*/*Aedes* abundance (right axis) at each subsite plotted against pH. **E**) Distribution of *Culex* abundance (left axis) and *Anopheles*/*Aedes* abundance (right axis) at each subsite plotted against relative turbidity.

### Incorporating rainfall data into model to predict *Anopheles* presence

Since each permanent/semipermanent site was visited twice during the sampling period, with at least one week separating each visit, differences in *Anopheles* larvae abundance between visits were analyzed. Significantly more *Anopheles* larvae were caught per subsite during the second visit of fountains (Two-tailed Mann-Whitney U test, *P* = 0.0169, U_(15)_ = 67.5) and wetlands (Two-tailed Mann-Whitney U test, *P*=0.0485, U_(22)_ = 187.0), while no significant difference in *Anopheles* larvae abundance between visits of artificial ponds and ditches was observed (**Fig. 4A**).

**Figure 4:**
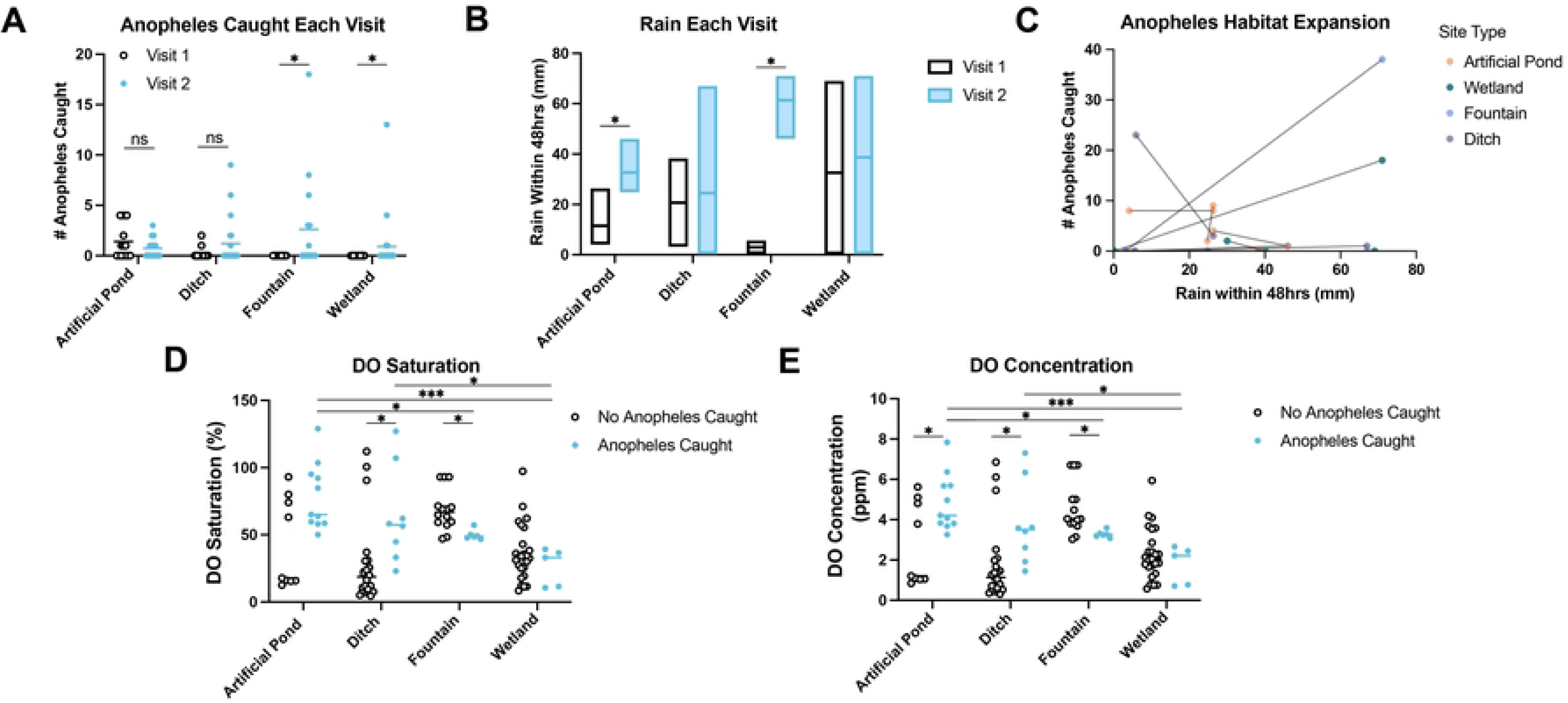
The effect of rainfall on *Anopheles* breeding site preferences. **A**) Distribution of Anopheles larvae abundance at subsites between site types and visits. **B**) Amount of rain within 48hrs of visiting each site, grouped by site types. Statistical significance determined using multiple Wilcoxon tests for paired data. **C**) Individual pairwise comparisons of Anopheles larvae abundance at each visit to each site. X axis shows the amount of rain within 48hrs of visiting that site. **D**) Distribution of percentage dissolved oxygen saturation across subsites of different site types, split into whether *Anopheles* larvae were caught. **E**) Distribution of dissolved oxygen concentrations across subsites of different site types, split into whether *Anopheles* larvae were caught. Statistical significance was determined using either a Mann-Whitney U Test, or Kruskal-Wallis Test with Dunn’s uncorrected step down tests, unless otherwise noted. For all pairwise comparisons: * *P*<0.05, \*\**P*<0.01, \*\*\**P*<0.001, \*\*\*\**P*<0.0001.

There was a significant difference in rainfall between visits of artificial ponds (Wilcoxon matched-pairs signed rank test, *P* = 0.00061, W = -108.0) and fountains (Wilcoxon matched-pairs signed rank test, *P*=0.000061, W = -120.0) (Fig. 4B).

However, because the visits for each type of permanent/semipermanent site did not occur on the same day (e.g. not all artificial ponds were sampled on the same day), comparisons of rainfall between individual sites were also made. Although wetlands had significant differences in *Anopheles* abundance between visits but nonsignificant differences in rainfall when subsites from multiple individual sites were analyzed together, one wetland had strong rain-associated increases in *Anopheles* larvae abundance that explain the significant results in panel A (**Fig. 4C**).

Because PCA analysis indicated an association between *Anopheles* breeding sites in Zanzibar City and higher dissolved oxygen levels, we hypothesized that rain- independent breeding sites might be preferred by *Anopheles* mosquitoes due to them having consistently higher levels of dissolved oxygen. Additionally, the semipermanent subsites sampled had significantly greater levels of dissolved oxygen (Welch’s T test, t_(94.69)_ = 6.912, *P*<0.0001), lower levels of salinity (Two-tailed Mann-Whitney U test, U_(84, 62)_ = 2053, *P* = 0.0146), and fewer predators present than permanent subsites (Χ^2^ = 5.227, df = 1, *P*=0.0222) (**Supplementary** Figure 2**, Supplementary Table 3**). The subsites from artificial ponds and ditches where *Anopheles* larvae were caught had significantly higher levels of dissolved oxygen than the subsites where no *Anopheles* larvae were caught (**Fig. 4D-E**) (**Supplementary Tables 4-5**). Subsites from fountains where *Anopheles* larvae were caught had significantly lower levels of dissolved oxygen than subsites from fountains without *Anopheles* larvae, and there was no significant difference in dissolved oxygen levels between either group of subsites from wetlands (**Supplementary Tables 4-5**). Regardless of the lack of association between dissolved oxygen levels and *Anopheles* presence in those site types, subsites from artificial ponds with *Anopheles* larvae had significantly higher levels of dissolved oxygen than subsites from fountains and wetlands with *Anopheles* larvae (**Fig. 4D-E**) (**Supplementary Tables 6-7**). Additionally, subsites from ditches with *Anopheles* larvae had significantly higher dissolved oxygen levels than subsites from wetlands with *Anopheles* larvae (**Supplementary Tables 6-7**).

### Regression Analysis

Given that the data suggest an interaction between dissolved oxygen levels (both saturation and concentration), rainfall within 48hrs, and/or site type, we fit a logistic regression model to all data from permanent/semipermanent sites and temporary sites with a binary outcome of “*Anopheles* caught” or “no *Anopheles* caught”. We included physical parameters that were significant in the original PCA regression analysis, as well as dissolved oxygen concentration and depth. We omitted data that were included in the initial PCA regression analysis, but for which dissolved oxygen data was unavailable. Dissolved oxygen concentration was included in the model instead of dissolved oxygen saturation because dissolved oxygen concentration was more significantly associated with *Anopheles* presence than saturation (as shown in Fig. 4D- E). Interaction terms included dissolved oxygen saturation and semipermanence, dissolved oxygen concentration and concrete, dissolved oxygen concentration and rainfall within 48hrs, concrete and rain within 48hrs, and natural and rain within 48hrs.

Significant predictors included the percentage of dips with *Aedes* larvae (lower odds of catching *Anopheles* larvae), increased depth (lower odds of catching *Anopheles* larvae), increased salinity (lower odds of catching *Anopheles* larvae), the presence of a predator in the quadrat (lower odds of catching *Anopheles* larvae), more rain within 48hrs (lower odds of catching *Anopheles* larvae) increased concentration dissolved oxygen in semipermanent subsites (lower odds of catching *Anopheles* larvae), increased concentration dissolved oxygen in concrete subsites (higher odds of catching *Anopheles* larvae), and increased concentration of dissolved oxygen after more rainfall within 48hrs (higher odds of catching *Anopheles* larvae) (**Table 1**). The area under the receiver operating characteristic (ROC) curve was 0.9034, and the positive predictive power was 65.7% (**Supplementary** Figure 3).

**Table 1:**
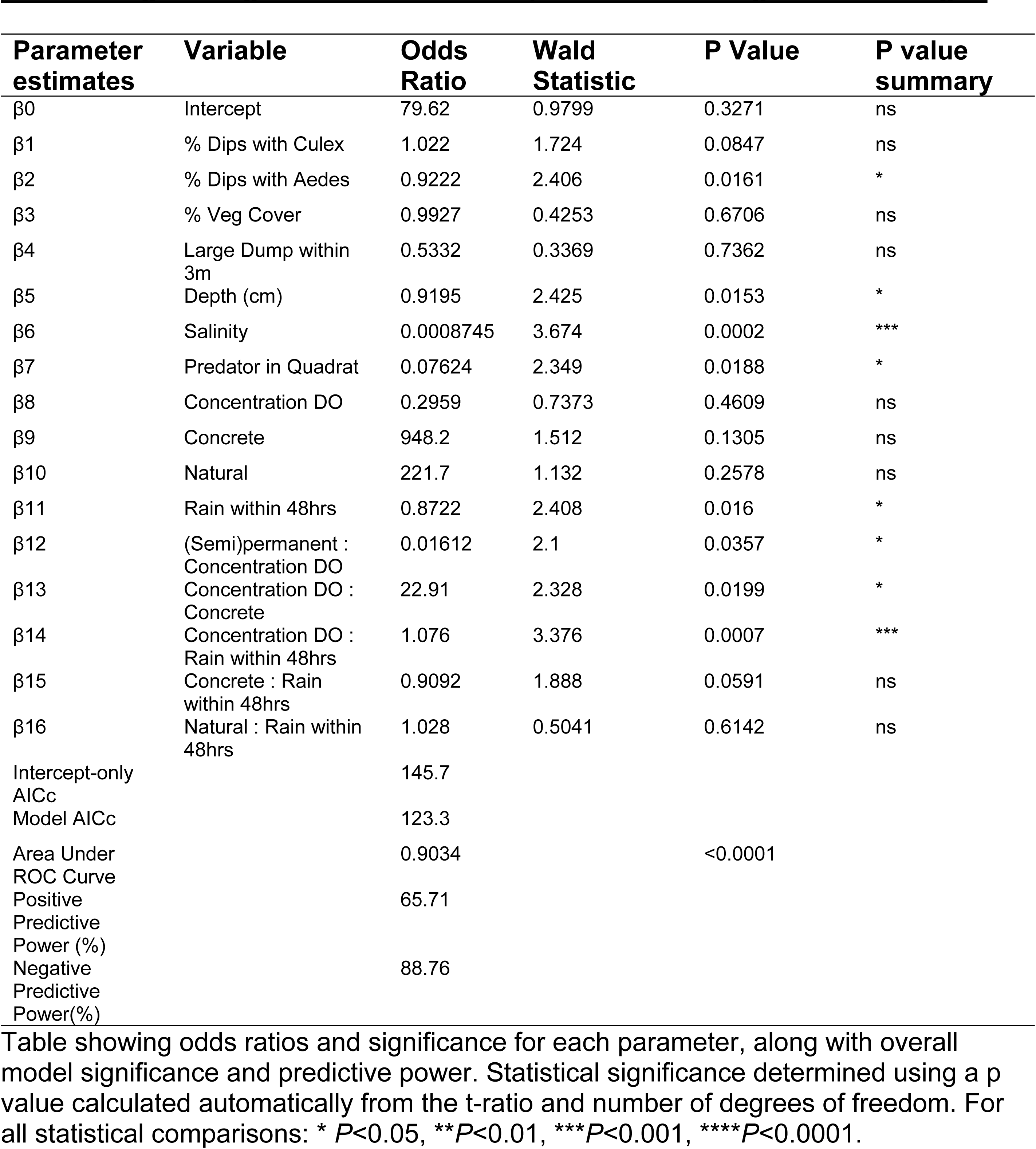
Logistic regression model of *Anopheles* larvae caught vs. not caught.

## DISCUSSION

In this study, we aimed to characterize the breeding sites of *Anopheles* mosquitoes during the rainy season in Zanzibar City. We found that a site being concrete, semipermanent, and having higher dissolved oxygen levels are significant predictors for *Anopheles* presence in Zanzibar City (regardless of rainfall). We also found that the interaction between higher rainfall levels and higher dissolved oxygen levels is a significant predictor for the presence of *Anopheles* larvae. These results suggest that *Anopheles* breeding sites in Zanzibar City tend to be concrete, semipermanent water bodies with high dissolved oxygen saturation regardless of heavy rainfall but can expand to natural semipermanent water bodies with high dissolved oxygen concentration after heavier rains.

Semipermanent, artificial sites are likely preferred over temporary breeding sites due to differences in developmental times of larvae from different mosquito genera.

*Anopheles* mosquitoes develop from eggs to adults in 10-14 days, while *Aedes* and *Culex* mosquitoes develop from eggs to adults in 7-10 days [36]. Since *Anopheles* larvae take longer to develop, they may either be outcompeted in temporary environments where *Aedes* mosquitoes are more suited to breed in, or the container is emptied too frequently for *Anopheles* larvae to develop [37]. This finding is consistent with studies in Ethiopia and elsewhere where more *Anopheles* larvae were found mostly in artificial, permanent/semipermanent sites [26,38–40]. However, this finding contradicts other studies where *Anopheles* larvae were found mostly in either natural or temporary environments, suggesting either a location and context-dependent habit to use semipermanent sites for breeding, or variation in the definition of “temporary” between studies [24,41,42]. Nevertheless, semipermanent sites may be preferred as *Anopheles* breeding sites over permanent sites because the semipermanent subsites sampled had higher levels of dissolved oxygen, fewer predators, and lower salinity than permanent sites (Supplementary Figure 2). Chemical qualities of water bodies influence *Anopheles* ability to breed due to impacts on nutrient availability, toxicity, selection for predators, algae growth, exposure to sunlight, and other factors [43–45]. According to our and others’ data, dissolved oxygen, low abundance of predators, and low salinity are associated with higher *Anopheles* larvae abundance in some locations but not in others [46,47]. While the reasons behind these differences in breeding site chemical characteristics are likely to be site-specific, permanent sites may have more artificial introduction of predators or competing species (as was clearly the case at an artificial pond outside of a hotel in Stone Town), have more accumulated pollution than the more frequently-drained semipermanent sites (which may be indicated by lower dissolved oxygen levels), or have higher levels of nutrients that support a naturally higher population of predators. Higher predator abundance (particularly fish and water bugs) has been associated with fewer *Anopheles* larvae in permanent sites in Ethiopia [45], which is consistent with our data, except semipermanent sites are used as breeding sites in Zanzibar City instead of temporary sites. Therefore, the data presented in this study support a location-dependent hypothesis for *Anopheles* breeding site preferences, where *Anopheles* prefer artificial, semipermanent breeding sites during the rainy season in Zanzibar City.

Although we observed statistically significant cohabitation between *Anopheles* and *Culex* larvae, this cohabitation was relatively uncommon, suggesting that each genus occupies distinct niches in Zanzibar City. This lack of common cohabitation is consistent with data from other studies, suggesting that *Culex* and *Anopheles* larvae compete for nutrients [45,48]. The site in our study with the most cohabitation between *Anopheles* and *Culex* larvae was a large, flooded wetland only after heavy rain, which could mean that the rain-associated expansion of this breeding site minimized the competition between mosquito genera. *Culex* larvae are well known to tolerate heavily polluted sites and are often able to avoid competition at these sites [49,50], while the tolerance of *Anopheles* larvae for polluted sites is more debated. While some study locations support the hypothesis that *Anopheles* primarily breed in clean sites [46,47], studies on mainland Africa have provided evidence of *Anopheles* adapting to more polluted environments [38,41,51,52]. With *Aedes* larvae dominating temporary breeding sites and *Culex* larvae dominating more polluted breeding sites with salinity levels above 0 ppt, it is possible that there is little pressure for *Anopheles* to use more polluted sites for breeding during the rainy season in Zanzibar City. However, as breeding site characteristics change during the dry season - when the semipermanent sites may not be available for breeding [39] - it is likely that *Anopheles* will need to find other breeding sites and may be forced into more polluted environments. Studies of *Anopheles* breeding sites in Zanzibar City during other seasons are, therefore, crucial for a complete understanding of breeding dynamics.

As high dissolved oxygen is an indicator of overall clean water, it is unclear from our data whether *Anopheles* larvae are found in sites with relatively higher dissolved oxygen concentrations/saturation because of the requirement for dissolved oxygen itself, or because of the lack of pollutants that negatively covary with dissolved oxygen levels. Data from experiments on *Culex* and *Aedes* larvae suggest that larvae from these genera can use dissolved oxygen for respiration when reaching the water surface to access atmospheric oxygen is difficult [53,54]. Although formal dissolved oxygen utilization experiments with *Anopheles* larvae are lacking, some authors suggest that dissolved oxygen might itself be especially important for *Anopheles* larvae development because *Anopheles* larvae lack breathing tubes [55]. Alternatively, as higher dissolved oxygen is an indicator of a less-polluted environment [56], the tendency for *Anopheles* breeding sites to have higher dissolved oxygen levels may be solely because those sites are less likely to have harmful pollutants and more likely to have the nutrients required for *Anopheles* development. Due to the lack of a strong, direct correlation between dissolved oxygen levels and *Anopheles* larvae abundance, the data from this study more strongly support the latter hypothesis (especially since the water surface was accessible to larvae in all breeding sites). However, more laboratory-based experimental studies are necessary to determine the role of dissolved oxygen in *Anopheles* larvae development.

Because this sampling period took place during the April rainy season, where consistent rain over many days is followed by days of intense sun, we hypothesized that rain might play a role in the changes in breeding site productivity between visits. Studies have shown that rainfall contributes to either the expansion of *Anopheles* breeding sites, or the flooding of breeding sites and increased mortality of *Anopheles* larvae [57,58].

However, these two outcomes are clearly dependent on the amount of rainfall and the specific shape of the site. Of all the sites sampled in this study, those most subjected to drainage coming into the site (and potentially bringing *Anopheles* larvae from other breeding sites) are those classified as drains, and the wetland located at the eastern extreme of the study area. Notably, a small number of *Anopheles* larvae were found at the wetland on the eastern extreme of the study site, but this occurred on the visit with less rain, so it is unlikely that the presence of *Anopheles* larvae at this site is dependent on runoff carrying the larvae into the wetland. All other sites were filled mostly by rainwater and did not accept runoff from other locations, so any increase in *Anopheles* larvae can be attributed to breeding at that site. Our data, therefore, suggest that higher rainfall levels are associated with more *Anopheles* larvae being found in wetlands and fountains, while artificial ponds and ditches serve as *Anopheles* breeding sites regardless of rain levels.

Limitations to this study include other potentially important parameters (e.g. algae cover) not being analyzed, limited geographical range, and a lack of species-level resolution of mosquito breeding habits. Differences in microbiota composition between sites were also not analyzed, which could also impact the ability for *Anopheles* larvae to develop [59]. While the purpose of this study was to analyze the *Anopheles* genus, as all *Anopheles* species in Zanzibar are capable of transmitting malaria, individual species have distinct breeding habits in other locations [25,60,61]. Determining which *Anopheles* species are occupying each type of breeding site in Zanzibar City would be useful for monitoring adaptations to more polluted breeding sites and the relative importance of each species in malaria and filarial disease transmission [31,41].

Additionally, larvicidal strategies tend to be more effective in areas where *Anopheles* adults are exophilic, as residual indoor spraying or bed nets are less effective in those areas [62]. Therefore, studies on the behavior of adult *Anopheles* mosquitoes in Zanzibar City are necessary and can also help further assess breeding site productivity [63]. Nonetheless, the short-term nature of the study provides evidence of rapid, rain-associated changes in *Anopheles* breeding site preferences in Zanzibar City, while also revealing sites that are likely to be more consistent breeding sites if they remain filled with water. The small geographical range included provides a targeted analysis of Zanzibar City-specific breeding habitats of *Anopheles* mosquitoes. Additionally, while our study did not include all potentially relevant parameters, the parameters measured are easy to monitor (e.g. more easily than site-specific microbiota) and can therefore be put into practice for larvicidal strategies targeted against *Anopheles*. This study can serve as a baseline for future longitudinal studies on *Anopheles* breeding sites in Zanzibar City, which could be used to develop a larvicidal strategy to combat persistent malaria in the Zanzibar archipelago.

Although larvicides are rarely used in Africa, [21,22] the data from our study suggest that semipermanent ponds or obstructed concrete ditches with high levels of dissolved oxygen saturation may be effective targets for larvicidal strategies in Zanzibar City. Additionally, preventing the flooding and expansion of wetland sites during rain by increasing drainage from those sites might also help prevent *Anopheles* from using these as breeding sites. Because mosquito larvae require stagnant water to breathe [64], larvae abundance in fountains could be minimized by ensuring that these fountains remain functional, and larvae abundance in drains could be minimized by ensuring that these drains remain free of stagnant water. The use of synthetic larvicides should be avoided, as their use can result in harmful, off-target effects to the environment, and mosquito larvae can develop resistance to the larvicides [65]. However, plant extracts may be a useful alternative, as they are more benign to the environment, more biodegradable, and contain a diverse array of larvicidal secondary compounds that minimize the risk of resistance [66,67]. Additionally, bacterial larvicides using larva-specific toxins from *Bacillus thuringiensis israelensis* or *Bacillus sphaericus* may be another safe and effective larvicidal strategy [68,69]. Introduction of predatory fish into artificial ponds could also prevent these locations from serving as *Anopheles* breeding sites [70]. Clearly, a variety of environmentally-friendly and sustainable larvicidal strategies exist that may aid in lowering the population of *Anopheles* mosquitoes in Zanzibar City.

## CONCLUSION

The incidence of malaria in Zanzibar City has remained between 1-2% since 2015, with a slight but steady increase due to continued import of *Plasmodium* spp. from other malaria endemic areas [6,7,9,10]. Therefore, targeting the resident population of *Anopheles* vector mosquitoes in the largest city of the Zanzibar archipelago is a crucial strategy to combating malaria on the islands. With pyrethroid resistance in adult *Anopheles* mosquitoes increasing on Unguja [71], larvicidal strategies could serve as an alternative strategy to decrease the population numbers of resident *Anopheles* mosquitoes. Although the geographic range of this study focused on the core of Zanzibar City, *Anopheles* larvae appeared to be evenly distributed between core Stone Town and the surrounding Zanzibar City. However, as most beautification projects and artificial, semipermanent structures are in the more tourist-trafficked area of core Stone Town, careful monitoring of these projects is necessary to prevent these concrete basins in parks from becoming nutrient and oxygen-rich breeding grounds for *Anopheles* mosquitoes.

Our study provides the first systematic survey of *Anopheles* breeding sites in Zanzibar City. Our findings that *Anopheles* prefer either semipermanent, concrete habitats with high dissolved oxygen levels, but can expand to sites with lower dissolved oxygen levels after heavy rains, suggest that introducing environmentally-friendly larvicides into artificial ponds, ensuring that fountains are functional, and preventing the flooding of wetlands may reduce the population of *Anopheles* mosquitoes during the rainy season.

## ACKNOWLEDGEMENTS

We would like to acknowledge and thank the faculty and staff at the School for International Training office in Zanzibar for providing logistical and material help for this project. Additionally, we thank the faculty at the University of Dar es Salaam Institute for Marine Science for lending us the water quality measurement devices.

## SUPPLEMENTARY FIGURES

**Supplementary Figure 1:**
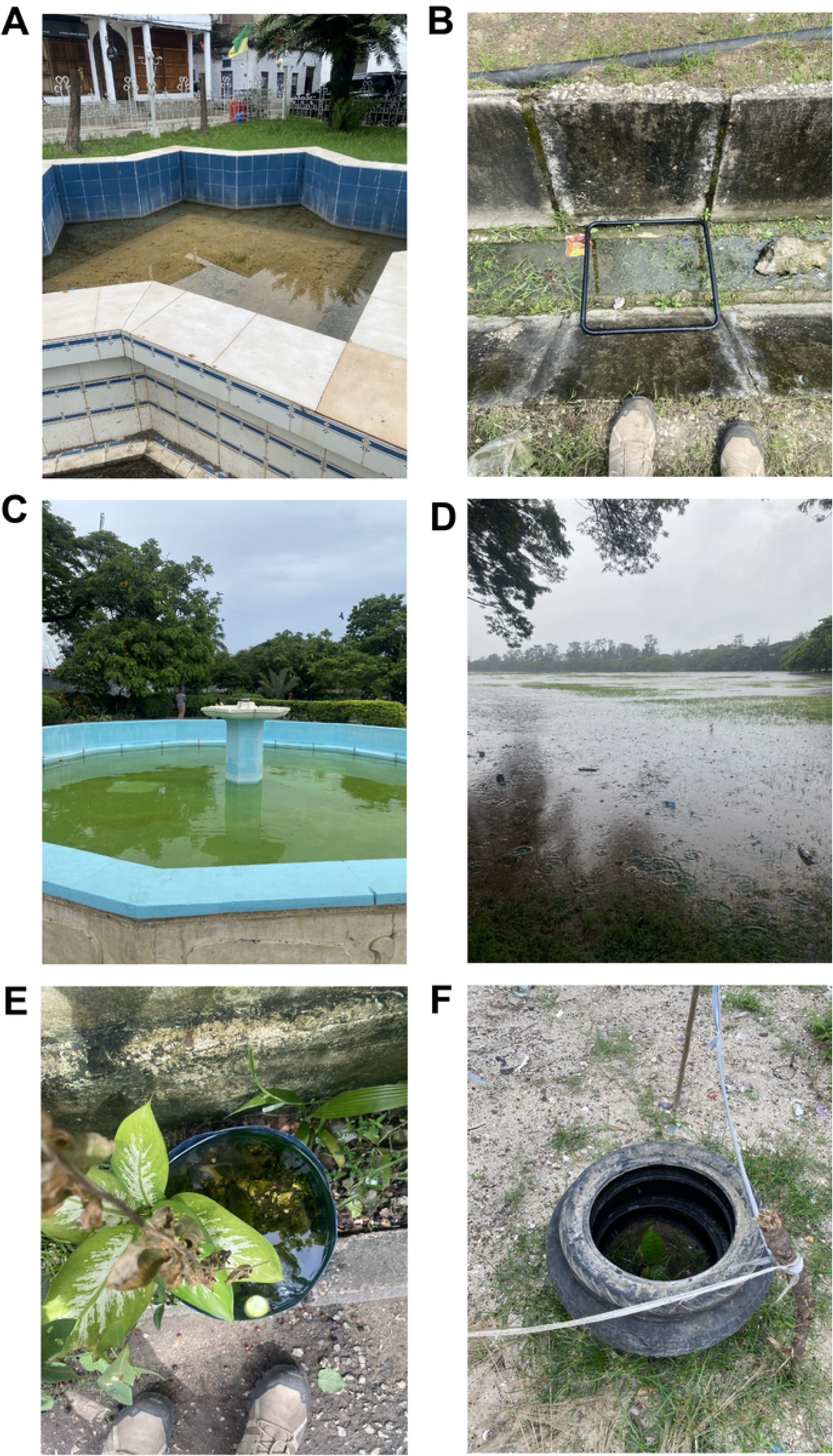
Photos of examples of each site type. **A**) Artificial pond. **B**) Ditch. **C**) Fountain. **D**) Wetland. **E**) Temporary water jug. **F**) Temporary tire.

**Supplementary Figure 2:**
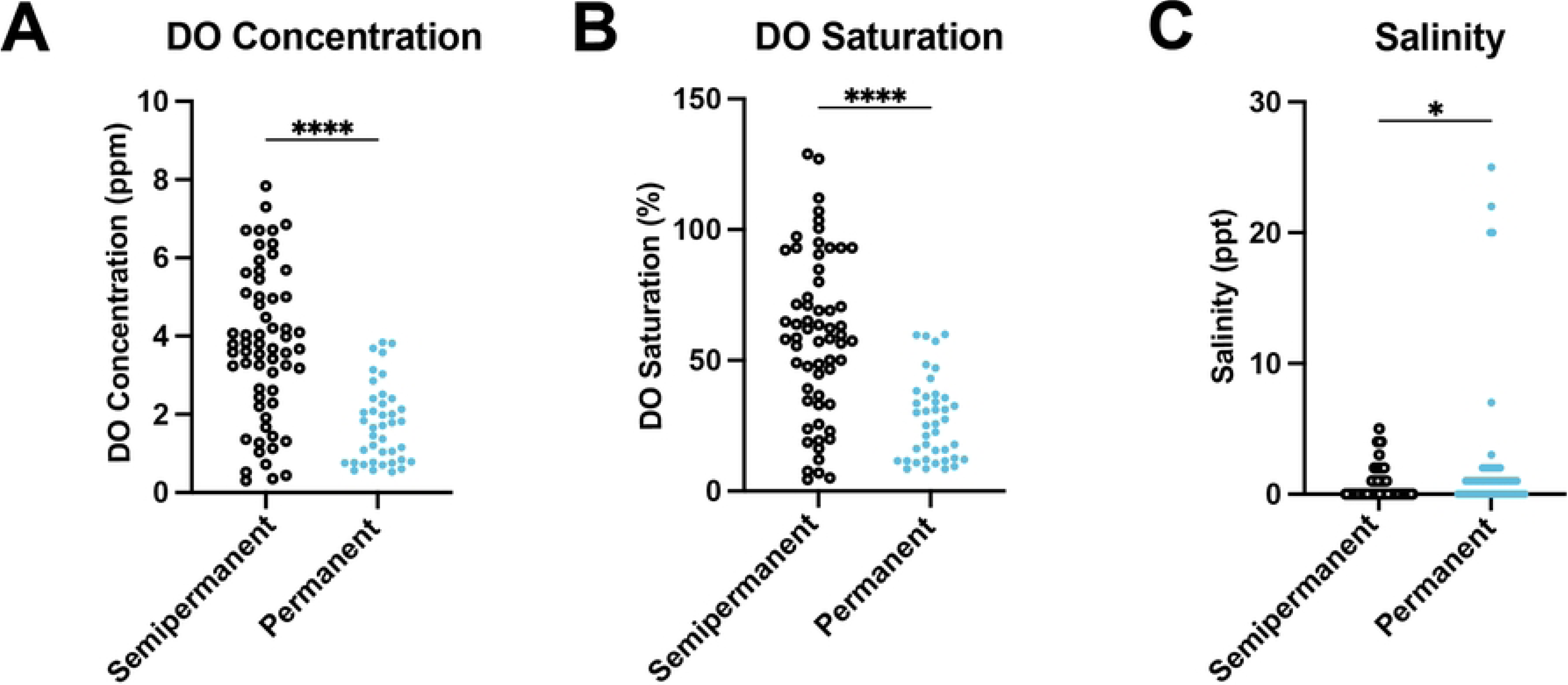
Differences in water quality parameters between semipermanent and permanent subsites. **A**) Dissolved oxygen concentrations in semipermanent and permanent subsites. **B**) Dissolved oxygen saturation percentages in semipermanent and permanent subsites. **C**) Salinity levels in semipermanent and permanent subsites. Statistical significance determined using Student’s unpaired T tests or Mann-Whitney tests for nonparametric data (salinity levels). For all pairwise comparisons: * *P*<0.05, \*\**P*<0.01, \*\*\**P*<0.001, \*\*\*\**P*<0.0001.

**Supplementary Figure 3:**
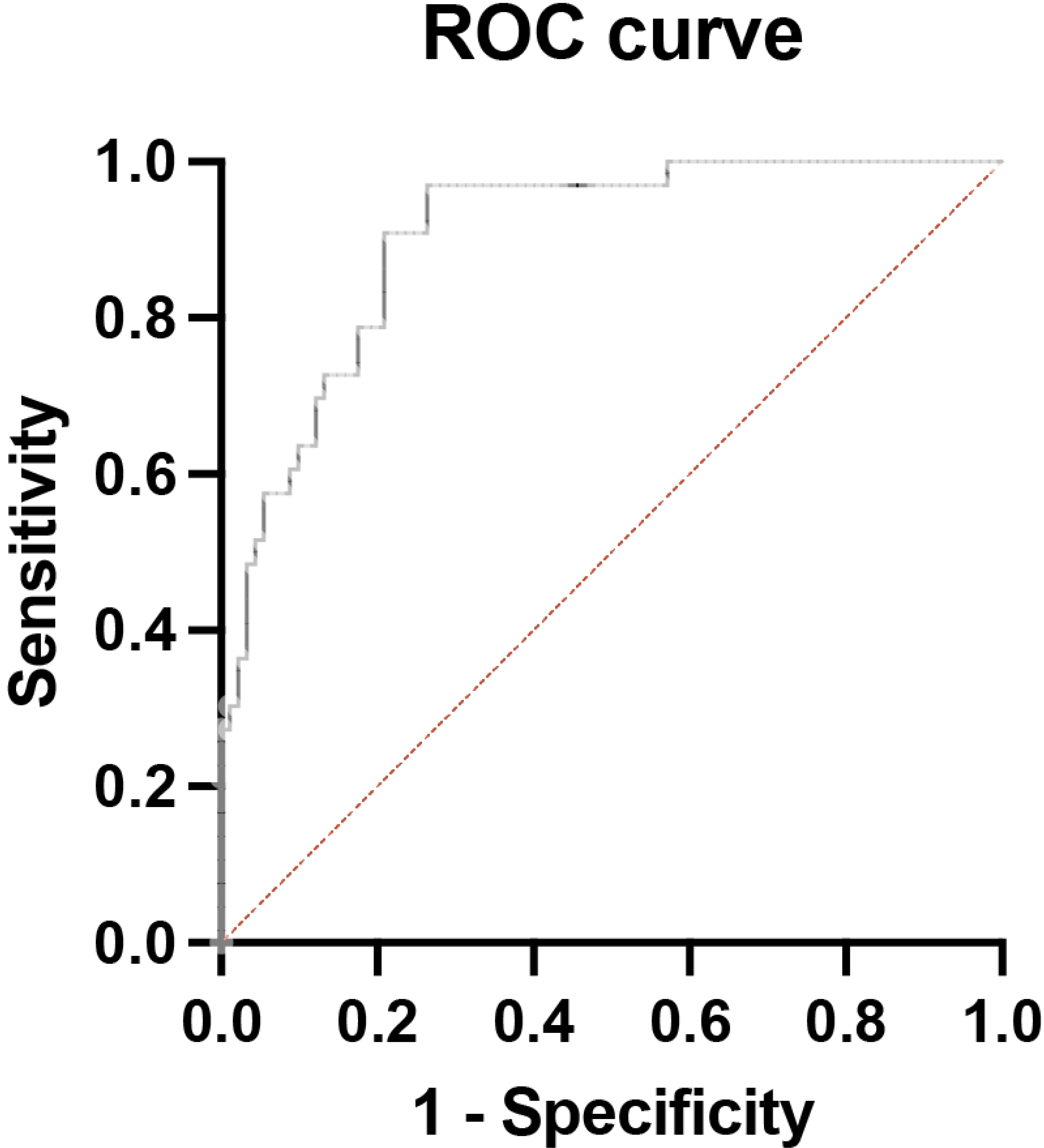
Receiver operator characteristic curve for the logistic regression model. Curve shows a deviation of the current model (black line) from a model with 50/50 odds of predicting *Anopheles* presence (red line).

## SUPPLEMENARY TABLES

**Supplementary Table 1:** Permanent/semipermanent site-level *Anopheles* larvae abundances.

| Site | Site Type | # Anopheles Visit 1 | # Anopheles Visit 2 |
| --- | --- | --- | --- |
| 1 | Artificial Pond | 8 | 8 |
| 2 | Artificial Pond | 0 | 0 |
| 3 | Artificial Pond | 4 | 1 |
| 4 | Artificial Pond | 9 | 2 |
| 5 | Ditch | 0 | 1 |
| 6 | Ditch | 0 | 0 |
| 7 | Ditch | 3 | 23 |
| 8 | Ditch | 0 | 0 |
| 9 | Fountain | 0 | 0 |
| 10 | Fountain | 0 | 1 |
| 11 | Fountain | 0 | 38 |
| 12 | Wetland | 0 | 0 |
| 13 | Wetland | 0 | 18 |
| 14 | Wetland | 0 | 2 |
| 15 | Wetland | 0 | 0 |
| 16 | Wetland | 0 | 0 |

**Supplementary Table 2:** Principal component regression results for physical parameters.

| Parameter estimates | Variable | Estimate | P value | P value summary |
| --- | --- | --- | --- | --- |
| $\beta_0$ | Intercept | 8.809 | 0.3014 | ns |
| $\beta_1$ | % Dips with Culex | -0.01914 | 0.0206 | * |
| $\beta_2$ | % Dips with Aedes | -0.02653 | 0.0223 | * |
| $\beta_3$ | Outdoor Temp (°C) | -0.1449 | 0.4581 | ns |
| $\beta_4$ | Water Temp (°C) | -0.1332 | 0.4283 | ns |
| $\beta_5$ | % Veg Cover | -0.02336 | 0.0349 | * |
| $\beta_6$ | Large Dump within 3m | -1.813 | 0.0399 | * |
| $\beta_7$ | Trash within 3m | -0.5038 | 0.2153 | ns |
| $\beta_8$ | Depth (cm) | 0.02395 | 0.1812 | ns |
| $\beta_9$ | Predator in Quadrat | 0.6232 | 0.0771 | ns |
| $\beta_{10}$ | Site Perimeter | 0.3423 | 0.1284 | ns |
| $\beta_{11}$ | (Semi)permanent | 2.394 | 0.0209 | * |
| $\beta_{12}$ | Artificial | 1.847 | 0.0311 | * |
| $\beta_{13}$ | Concrete | 2.874 | 0.0034 | ** |
| $\beta_{14}$ | Plastic/Rubber | -2.425 | 0.0193 | * |
| $\beta_{15}$ | Natural | -1.847 | 0.0311 | * |
| $\beta_{16}$ | Visit | 1.393 | 0.06 | ns |
| $\beta_{17}$ | Rain within 48hrs | 0.01728 | 0.3501 | ns |
| Regression ANOVA<br>F Statistic |  | 3.288 | 0.0308 | * |

All slope estimates are presented in terms of the parameters, not in terms of principal components. Statistical significance for each parameter determined automatically using a T-distribution. Overall significance for the regression determined using ANOVA. For all statistical comparisons: \* $p < 0.05$ , \*\* $p < 0.01$ , \*\*\* $p < 0.001$ , \*\*\*\* $p < 0.0001$ .

**Supplementary Table 3:**
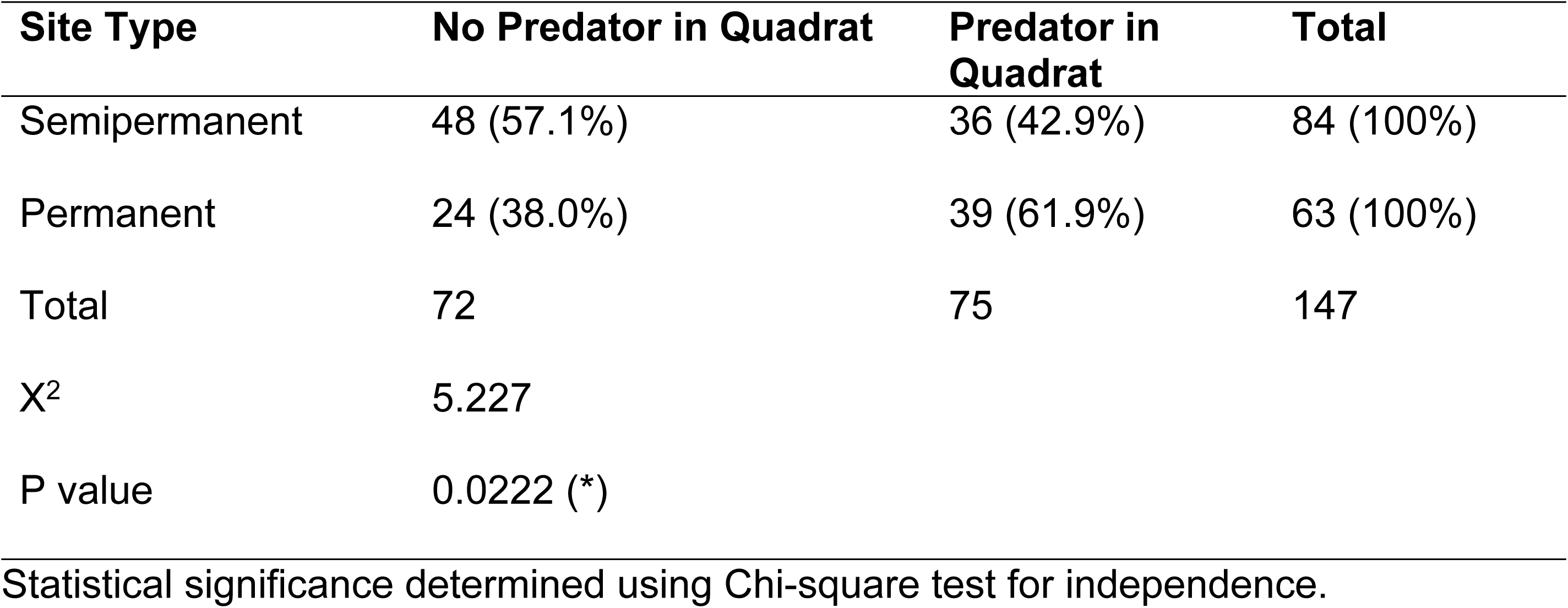
Differences in predator abundance between semipermanent and permanent subsites.

| Site Type | No Predator in Quadrat | Predator in Quadrat | Total |
| --- | --- | --- | --- |
| Semipermanent | 48 (57.1%) | 36 (42.9%) | 84 (100%) |
| Permanent | 24 (38.0%) | 39 (61.9%) | 63 (100%) |
| Total | 72 | 75 | 147 |
| X<sup>2</sup> | 5.227 |  |  |
| P value | 0.0222 (*) |  |  |

Statistical significance determined using Chi-square test for independence.

**Supplementary Table 4:**
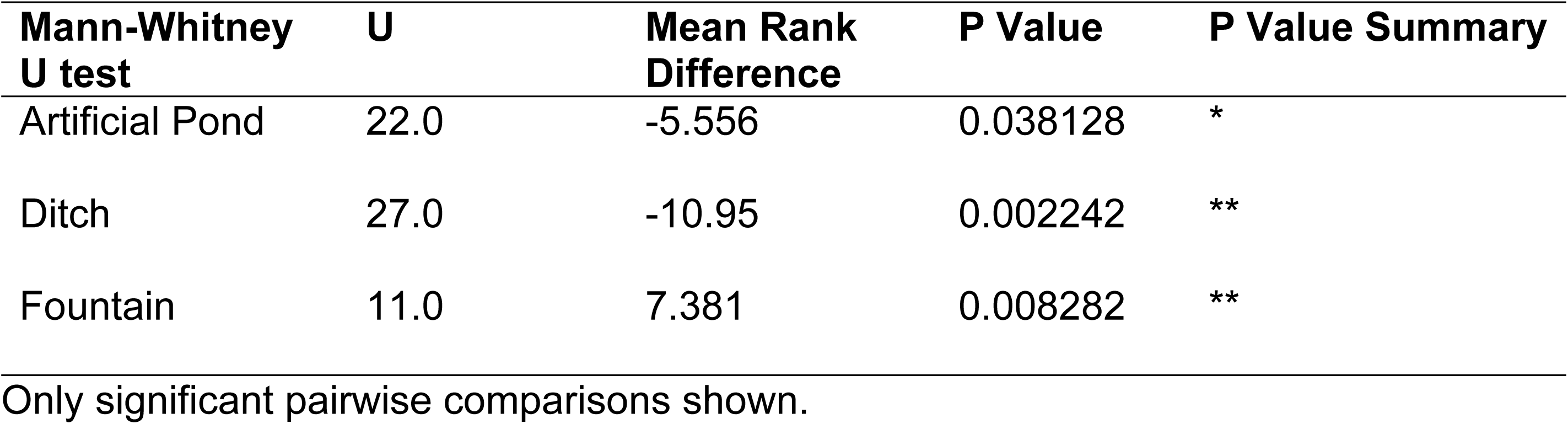
Two-tailed Mann-Whitney U test results comparing oxygen concentration between subsites with and without *Anopheles* larvae, split by site type.

| Mann-Whitney U test | U | Mean Rank Difference | P Value | P Value Summary |
| --- | --- | --- | --- | --- |
| Artificial Pond | 22.0 | -5.556 | 0.038128 | * |
| Ditch | 27.0 | -10.95 | 0.002242 | ** |
| Fountain | 11.0 | 7.381 | 0.008282 | ** |

Only significant pairwise comparisons shown.

**Supplementary Table 5:** Two-tailed Mann-Whitney U test results comparing oxygen saturation between subsites with and without *Anopheles* larvae, split by site type.

| Mann-Whitney U test | U | Mean Rank Difference | P Value | P Value Summary |
| --- | --- | --- | --- | --- |
| Ditch | 25.0 | -11.29 | 0.001529 | ** |
| Fountain | 9.00 | 7.857 | 0.004438 | ** |

Only significant pairwise comparisons shown.

**Supplementary Table 6:** Kruskal-Wallace test results comparing dissolved oxygen concentration at subsites with Anopheles between different site types.

| Kruskal-Wallace Test | H = 16.18 | Groups = 4 | P = 0.001 |  |
| --- | --- | --- | --- | --- |
| **Dunn's Test for Multiple Comparisons** | **Z** | **Mean Rank Diff.** | **P Value** | **P Value Summary** |
| Artificial Pond vs. Fountain | 2.326 | 10.39 | 0.0200 | * |
| Artificial Pond vs. Wetland | 3.860 | 18.33 | 0.0001 | *** |
| Ditch vs. Wetland | 2.087 | 10.48 | 0.0369 | * |

Only significant pairwise comparisons shown.

**Supplementary Table 7:** Kruskal-Wallace test results comparing dissolved oxygen saturation levels at different site types.

| Kruskal-Wallace Test | H = 16.32 | Groups = 4 | P = 0.001 |  |
| --- | --- | --- | --- | --- |
| **Dunn's Test for Multiple Comparisons** | **Z** | **Mean Rank Diff.** | **P Value** | **P Value Summary** |
| Artificial Pond vs. Fountain | 2.349 | 10.49 | 0.0188 | * |
| Artificial Pond vs. Wetland | 3.877 | 18.41 | 0.0001 | *** |
| Ditch vs. Wetland | 2.366 | 11.88 | 0.018 | * |

Only significant pairwise comparisons shown.

